# Using Topological Data Analysis and RRT to Investigate Protein Conformational Spaces

**DOI:** 10.1101/2021.08.16.456547

**Authors:** Ramin Dehghanpoor, Fatemeh Afrasiabi, Nurit Haspel

## Abstract

An essential step to understanding how different functionalities of proteins work is to explore their conformational space. However, because of the fleeting nature of conformational changes in proteins, investigating protein conformational spaces is a challenging task to do experimentally. Nonetheless, computational methods have shown to be practical to explore these conformational pathways. In this work, we use Topological Data Analysis (TDA) methods to evaluate our previously introduced algorithm called RRTMC, that uses a combination of Rapidly-exploring Random Trees algorithm and Monte Carlo criteria to explore these pathways. TDA is used to identify the intermediate conformations that are generated the most by RRTMC and examine how close they are to existing known intermediate conformations. We concluded that the intermediate conformations generated by RRTMC are close to existing experimental data and that TDA can be a helpful tool to analyze protein conformation sampling methods.

## 1 Introduction

Proteins perform vital functions, such as catalyzing metabolic reactions, regulating cell signaling, DNA replication, and so forth. They often go through large-scale conformational transitions to perform these functionalities. To understand and interpret protein functionalities, we need to understand their structures, their conformational spaces, and the pathways they take when going through these transitions. Nevertheless, the conformational space of proteins is complex and high dimensional, and hence studying it becomes a challenging task. Several attempts have been made to characterize protein conformational pathways and find intermediate structures created during these transitions. Experimental methods such as X-ray crystallography [1], NMR [2], and cryo-Electron Microscopy [3] have been used. However, because of the rapid dynamics of proteins, these changes happen in microseconds, and experimental methods cannot capture all of them. For this reason, there has been an increasing interest in computational methods to simulate and explore these pathways. Computationally, it is still a challenging problem to simulate these conformational changes because physics-based simulations take considerable time. Molecular Dynamics (MD) simulations can simulate these transitions at the microsecond level, but they are still not fast enough to capture transitions that take place in larger timescales [4]. Therefore, researches have come up with variants of MD simulations such as Steered MD [5], Replica Exchange MD [6], and Targeted MD [7]. A number of other methods attempt to explore the geometrically constrained conformational space by using the degrees of freedom (dihedral angles) to perturb the protein structure. A few instances of these search methods are Robotic Motion Planning[8], elastic network model [9], and normal mode analysis [10]. The advantage of these search methods is that they run much faster than MD simulations. However, they may result in pathways that are not physically feasible. We will review this issue more in section 1.1.

It is important to do an accurate sampling of the conformational space to identify intermediate conformations that correspond to local energy minima and are close to actual intermediates. Thus, there is a need to extract biologically relevant conformations and filter out implausible ones. To do so, biological data can be incorporated to guide the search to produce meaningful conformations. For example, rigidity information about the protein molecules [11] can provide useful knowledge about the rigid parts of the molecule, so that only flexible parts can be manipulated. This information can be gathered experimentally, from coevolution data, from computational analysis, or by using rigidity analysis [12, 11]. In section 1.1, we provide an overview of our previous work, and in this article, in section 2, we evaluate how close our generated intermediate conformations are to experimentally known intermediates.

### 1.1 Overview of RRT* Search and TDA

#### RRT* Search

The Rapidly exploring Random Tree (RRT) algorithm [13] is used to rapidly explore constrained spaces. It builds a tree rooted at the start configuration and grows by using random samples from the exploration space and connecting them to the closest vertex in the tree. We used this algorithm in our previous efforts, [14, 15] to explore the conformational space of proteins. Here is a quick review of the basic algorithm: At every iteration, the algorithm generates a random conformation *q_rand_*, finds the closest node to *q_rand_* in the tree, and tries to create a path from the closet node found to the *q_rand_*. If *q_rand_* has high energy, it repeats the same steps of generating the random conformations as long as the energy is below a threshold. The step is repeated until the new conformation is close enough to the goal conformation or the maximum number of iterations is reached. A detailed explanation of RRT and how it is used to explore the conformational space of proteins can be found in [15].

RRT explores the search space rapidly, but given that it randomly expands to everywhere in the search space, it may not converge fast enough for complex spaces like protein conformations. For this reason, we added a guided selection step that biases the search (with a probability of 1/3) towards a conformation among the top *k* conformations with the smallest least RMSD (lRMSD with respect to the goal conformation) instead of selecting from the entire pool of conformations. *k* is a user-defined input, and in our work, it is set to 20.

Additionally, as a result of the randomness in RRT, the produced paths could be uneven and rugged. To tackle these issues, the RRT* algorithm [16] optimizes paths by proposing a heuristic cost function to find less costly (smoother) parents for every new node that is added to the tree and changing parents if needed.

#### Rigidity Analysis

Proteins can be modeled as mechanical structures made up of rigid and flexible parts [11]. Rigid parts denote groups of atoms that are more likely to stay together during conformational changes. Taking the rigidity analysis information into account helps us gain insights into potential motions of proteins and select non-rigid (flexible) parts for manipulation. We used the Kinari software [11] in our previous work to incorporate rigidity analysis information to better guide our search algorithm. It implements a faster and more robust variation of the pebble game algorithm to extract rigidity information about the molecules without having to run extensive MD simulations.

#### TDA and the Mapper Algorithm

Topological Data Analysis (TDA) has been employed in recent years to analyze complex high-dimensional datasets and extract valuable information and insights [17]. TDA has been extensively used in biomedical research [18, 19]. By using TDA methods, researchers have found a way to understand the structures and the underlying shapes of high-dimensional datasets. Other methods have been introduced before, such as Principal Component Analysis (PCA), Multidimensional Scaling (MDS), etc. However, these methods may not catch all large and small-scale patterns existing in the data as well as TDA.

The Mapper algorithm [20] is an unsupervised learning TDA algorithm that utilizes a combination of dimensionality reduction, clustering, and graph networking methods to provide a graph (called the Mapper graph) that summarizes the structure of a dataset. The general idea of the Mapper algorithm is as follows: Suppose that we can divide our data by a measure of similarity into groups (determined based on the data) and represent each group by a node. Next, let us present the relationships between two groups through edge connections. This results in a simple graph that can visualize the structure of our complex data provided that we choose appropriate measures of similarity and clustering techniques. Fig. 1 shows an example of the Mapper algorithm where the data points of a 2D torus (A) are taken as input. In our work, the data points are protein conformations.

**Figure 1:**
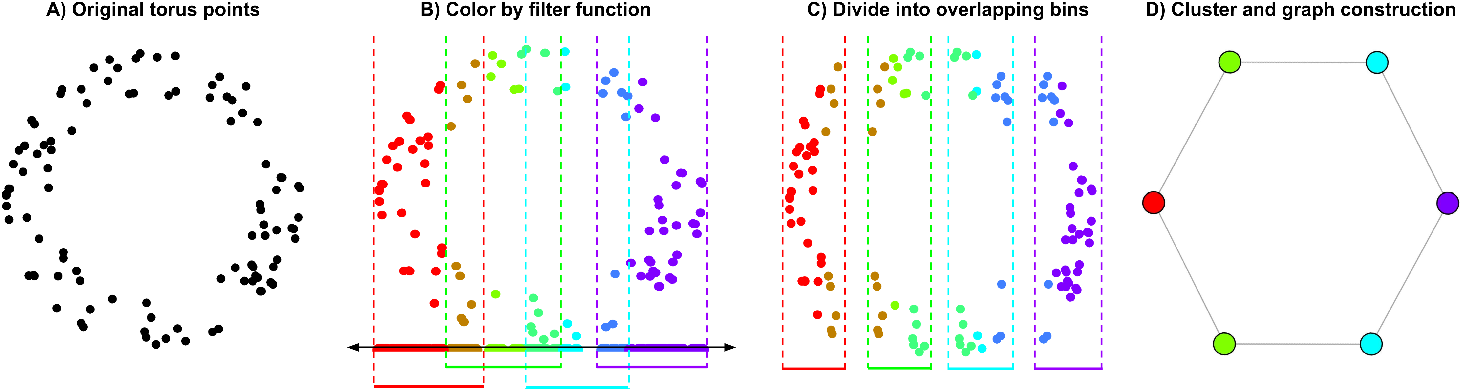
Schematic illustration of the Mapper algorithm run on the data points of a 2D torus.

A *filter function* is used to project the high-dimensional data points to a low-dimensional space where local relationships are preserved. In the torus example, the *filter function f*: ℓ^2^ → ℓ, (*x, y*) → *x* is used to project each data point to the *x* axis. The Mapper then creates a cover of the projected space as a set of partially overlapping bins (intervals). In Fig. 1 (B), the overlapping bins are shown in different colors, and in (C) they are shown separately. Note that *dark gold* colored data points, for instance, are the points that fall into both *red* and *green* colored bins. Next, data points within each cover are clustered based on their distances in the original space. Finally, Mapper constructs a graph representing the data where clusters are vertices and there is an edge between any two clusters that have a data point in common (D).

The advantage of the Mapper algorithm over other dimensionality reduction methods is that the traditional methods may suffer from projection loss. This happens when data points (protein conformations in our work) that are apart in high dimensions may be projected close together in the reduced lower-dimensional space. However, in Mapper, the distance between the original data points is used so that the covering in the projected space is pulled back to the original space, and then the clustering happens here. This is the advantage since in Mapper, the substructures that exist in the original high-dimensional space are found and used in covering and finding more relevant clusters. Mapper can visualize the complex original space in a compressed graph representation where the relationships between the original data points are shown in clusters and the edges between the clusters. Another fascinating feature of Mapper, which we use in section 3, is that when clusters and connected components are generated, it allows investigating why these data points are interconnected. For instance, by coloring the network by a quantity of interest (such as the energy of the conformations in our case), we can explain the various groups that have appeared and the distinctions between them.

#### Previous Work

In our previous work [14], we focused specifically on rigidity analysis information and extracted the flexible residues. We used this information to lead the search by using flexible hinges to manipulate. This approach helped RRTMC choose among a limited number of dihedral angles to rotate in order to perform the conformational transition. Since the rigidity properties of proteins may change when they go through large-scale conformational changes, we performed updated rigidity analysis during the run. To balance efficiency and accuracy, we found that the optimal point to recalculate the rigidity analysis was when there was a change of > 2*Å* from the last point of recalculation.

#### This Contribution

In this study, we do an evaluation of RRTMC using topological data analysis. It is based in part on previous work, but the novelty includes running RRTMC on more protein molecules with the integration of rigidity analysis, reducing the dimensionality of the conformational pathway data, and using TDA to construct a simple graph that represents the highly frequent intermediate structures. By using PCA, we reduce the dimensionality of our data, and then, by using the TDA Mapper algorithm, we construct a graph of our reduced dataset. We use about 200 runs of one of our proteins from our test case. We then extract the intermediate conformations that appear the most in the pathways produced by RRTMC and compare them with known experimental results. We used Adenylate Kinase (AdK) as input to the Mapper algorithm since it has experimentally known intermediate structures. The purpose of this paper is to evaluate RRTMC and also to demonstrate that TDA can be useful in assessing protein conformation simulation methods.

## 2 Materials and Methods

### 2.1 Data Collection and Preprocessing

In this contribution, our primary goal is to evaluate whether RRTMC produces conformations that are likely to represent intermediate conformations that exist in nature. To achieve this goal, we first reduce the dimensionality of the conformations that RRTMC generates and then use the Mapper algorithm to extract conformations that appear frequently and then compare them with known conformations of AdK. Experimentally discovered intermediate conformations (PDB files) are not available for many proteins, so we chose AdK to do our first evaluation. We intend to find more available intermediate structures in our future work to perform better evaluations. Because of the randomness in the nature of robotic motion planning algorithms, we run the algorithm several times to make sure we have a reliable coverage of the conformational paths produced by our algorithm (RRTMC).

We ran the algorithm on each test case at least 150 times, and for AdK we did about 450 times (both 1*AKE* → 4*AKE* and 4*AKE* → 1*AKE*). In RRTMC, iterative sampling of new conformations continues up to the point that the score of the newly generated conformation (*lRMSD* to the goal conformation) is below a preset threshold or the minimum score achieved by generated conformations does not decrease for 500 iterations. The preset *lRMSD* threshold is between 1.7*Å* and 4*Å*, based on the protein size and pathway difficulty. In order to find the best threshold that would be a tradeoff between time and how close our conformations are to the goal conformation, we tried different thresholds for each of our test cases. Fig. 2 shows the result for some of our examples including adenylate kinase (AdK), calmodulin (CaM), DNA, and lysine/arginine/ornithine-binding (LAO). We also ran RRTMC on cyanovirin-N (CVN) and ribose-binding protein (RBP) that were mentioned in our previous work. This plot shows a realistic lower bound for our test case proteins. Realistically, *lRMSD* values below 1.5-2Å are hard to achieve due to sampling errors and variations in proteins. Each protein was represented using the backbone atoms (N, C-alpha, C, O) and C-beta.

**Figure 2:**
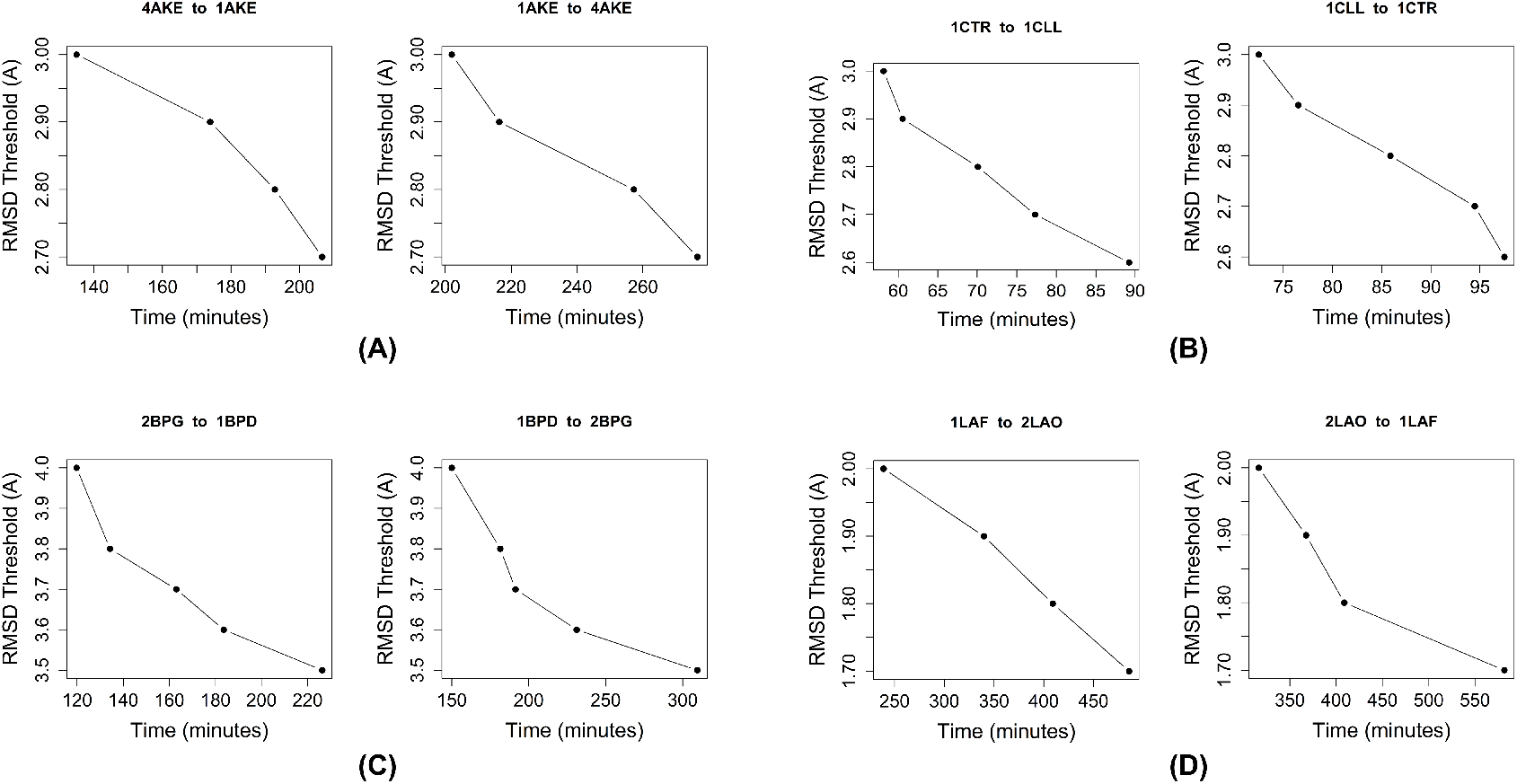
lRMSD threshold versus runtime average in minutes for (A) AdK, (B) CaM, (C) DNA, and (D) LAO.

To run the Mapper on our test case protein (AdK), in this work we only use runs of one direction; 4*AKE* → 1*AKE* (open conformation to closed conformation). Each path created an average of ten conformations from the start conformation (4*AKE*) to the goal conformation (1*AKE*). This data consist of a total of 1,415 intermediate conformations that form our highdimensional dataset. Each path is stored as a numeric matrix of *x, y, z* coordinates with one conformation per row and a Cartesian coordinate per column. That means, for instance, for a path of 20 intermediate conformations going from 4AKE to 1AKE, we get a matrix of 20 rows and 3 * 1050 columns (since AdK has 1,050 backbone atoms). Adding all the paths gives us a matrix of 1415 × 3150.

### 2.2 Using Mapper for Evaluation

We used Mapper implementation by Paul Pearson et al. [21] (accessed on 8 October 2021), which is an **R** package for using discrete Morse theory to analyze data sets.

Our 3-dimensional data, which are the results of running PCA on our high-dimensional data (Refer to 2.1), are divided into partially overlapping bins as described above. We use the default clustering method, single linkage, to cluster the data points in each bin. Finally, a graph is created from the set of clusters (graph vertices) and whether they have data points in common (graph edges). The choice of parameters is discussed in section 3.1. The resulting graph is shown in Fig. 3. The size of each vertex in the figure is proportional to the number of data points in the cluster presented by that vertex. We use different strategies to color the vertices of the graph to gain useful information. In Fig. 3 (A), we color the graph according to the average energy values of conformations that reside in the same cluster. In (B), to get to at least 90% of the total variance, we use principal component four (PC4) to color the vertices. For each cluster represented by a vertex, we calculate the average value of PC4 and color that vertex accordingly. In (C), we colored every vertex by their lRMSD regarding the goal conformation. Each Mapper graph in Fig. 3 is discussed extensively in section 3.1.

**Figure 3:**
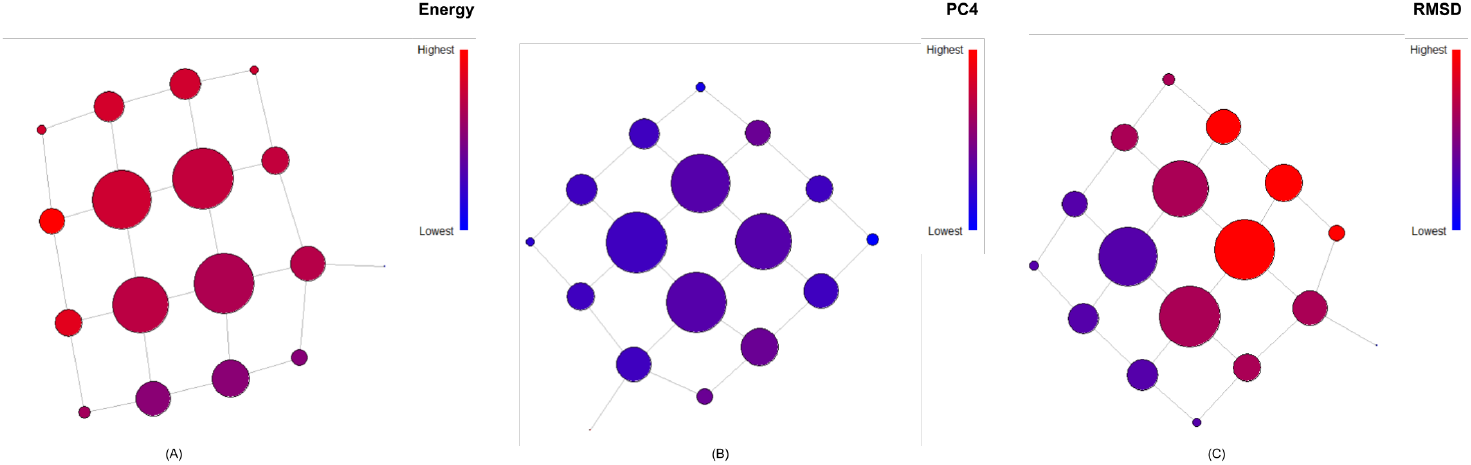
Mapper graph for AdK. Vertices are colored by average values of (A) energy, (B) PC4, and (C) lRMSD to the goal conformation

## 3 Results

### 3.1 Adenylate Kinase (AdK)

AdK is a monomeric phosphotransferase enzyme that catalyzes the reversible transfers of a phosphoryl group from ATP to AMP. AdK contains 214 amino acids and has three main domains: the CORE, the ATP binding domain called the LID, and the NMP binding domain. It is an interesting protein to study as far as conformational changes because it’s a signal-transducing protein, and there should be a balance between its conformations for it to regulate protein activity.

We used the following parameters to create a simple graph representing the structure of the data (Refer to section 1.1 for an overview of Mapper):

- distance-matrix: The distances between points. We used the Euclidean distances.
- filter-value: The vector of the projection of the data points generated using the filter function. We used PCA as our filter function to reduce and project the high-dimensional data to three dimensions. We used PC1, PC2, and PC3.
- num-intervals: Number of intervals in the co-domain of the filter function (regular grid over the filter range).
- percent-overlap: Percentage of overlap between the intervals.
- num-bins-when-clustering: controls the number of clusters for each inverse image (number of intervals of the histogram of edge lengths in the single linkage clustering algorithm)

Since mapper is an unsupervised learning algorithm and in unsupervised learning algorithms, the user usually selects the parameters, we applied a grid search to find the best parameters. We aimed to find the best parameters in our grid search so that not all data points would be clustered into a few big clusters, and at the same time, they wouldn’t be clustered into several very small clusters that don’t correctly demonstrate the relevant relationships between them. For our grid search, we used the following parameter ranges: {3, 4, 5, 6} for the number of intervals, {35, 40, 45, 50} for the overlap percentage, and {10, 11, 12, 13, 14, 15} for the number of bins when clustering. For these ranges, we found that for AdK, 4 for the number of intervals, 35 for the percentage of overlap, and 13 for the number of bins work the best. We chose these parameters this way to avoid either having all the data points (conformations) be clustered into one big cluster or having a large number of clusters with only a few data points in each of them.

In Fig. 3, the resulting graph has 17 vertices showing the clusters that contain all of our data points. Some points may be in more than one cluster (because of the overlapping bins), and where that happens, there is an edge connecting the two clusters. The size of each vertex is proportional to the number of points that reside in that cluster. That means that we have four clusters that contain most of our data points, where the largest one contains 445 conformations. These are the conformations that have been generated the most, and they are more expected to be close to highly populated regions. Small clusters mean that the conformations in those clusters are not likely to occur – for example, due to having high energy, and should probably be disregarded. Fig. 3 (A) shows the graph colored by the energy value. In our previous work, to calculate the energy value of our conformations, we used the following function [22]:

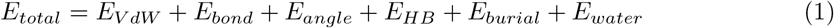

*E_VdW_, E_bond_*, and *E_angle_* are used from the *AMBER ff* 03 force field [23], where *E_VdW_* is the Lennard Jones potential [24]. However, it is adapted in order to tolerate soft collisions between atoms. We used these terms in our previous work in RRTMC to correct the clashes and deformities in the structure. Hydrogen bonds, *E_HB_*, burial *E_burial_*, and water-mediated interaction, *E_water_* terms are adapted from [22]. The cluster with the highest energy value has an average energy of 4251*kJ/mol*, and the one with the lowest energy value has an average value of –365*kJ/mol*. Many of the conformations have energy values between these numbers, which may correspond to local energy minima. Notice that energy values may be high due to the coarse-grained representation of the molecule.

In Fig. 3 (C), we can see the vertex colors relative to their lRMSD values from the goal conformation (the closed conformation, PDB: 1*AKE*). These colors are relative to each other, and it does not mean that the highest lRMSD is far from the goal conformation. The highest lRMSD to the goal conformation for the biggest clusters is less than 5.5*Å* (the start-goal lRMSD is approximately 7Å). The reason is that RRT is faster in the first steps, and it gets slower by getting closer to the end conformation. From the Mapper graph, we obtained the nine biggest clusters (since the average number of conformations in the simulated paths is ten and the first one is the start conformation), calculated the average coordinates of all the conformations in each cluster as the conformation representative of that cluster, and then for each representative conformation, calculated their lRMSD with respect to the goal conformation.

Using the first three principal components covered 78% of the total variance. By using PC4, we get close to 90%. It is not possible to use 4 vectors for the TDA mapper. Therefore, we used PC4 to color the vertices (Fig. 3 (B)). Fig. 4 shows the superimposition of the intermediate conformations on the path from 4AKE (shown in red) to 1AKE (shown in blue) that were extracted by Mapper. We used the biggest clusters, averaged the coordinates of the conformations in each cluster, and used these coordinates to create the intermediate conformations shown in the figure. As seen, the conformations follow a hinge motion.

**Figure 4:**
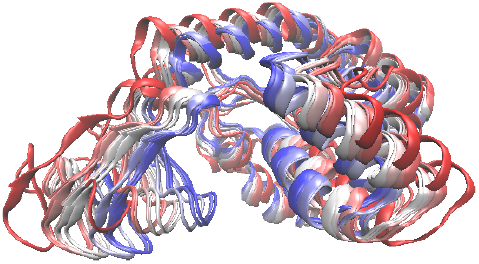
Superimposition of the intermediate conformations produced by Mapper for AdK - 4AKE (red) → 1AKE (blue)

### 3.2 Comparison to Experimental Results

As mentioned in section 1, experimentally known intermediate conformations are hard to find. Because of that, it is elusive to do experimental validation of computational methods. We used AdK, for which there exist several intermediate experimental structures [25]. We used our highly populated conformations, created by taking the average coordinates of conformations in each of the nine biggest clusters, and calculated their RSMDs with respect to the start, intermediate, and goal conformations. Table 1 shows that we could find clusters that the conformation created by their average coordinates are as close as 2.6*Å* to experimentally known conformations. We will explore other clustering methods in the future to determine whether we can get clusters that are even closer to experimental results.

**Table 1:** Comparison of Mapper-generated intermediates clusters to known intermediates

| protein | PDB ID<br>of known<br>intermediates | IRMSD ($\text{\AA}$) to closest | | Number of<br>conformations with<br>IRMSD $< 3\text{\AA}$ |
| --- | --- | --- | --- | --- |
|  |  | **mapper cluster**<br>**representative** | generated<br>conformation |  |
| AdK | 2AK2 | **3.7** | 3.4 | 0 |
|  | 1AK2 | **3.8** | 3.4 | 0 |
|  | 1DVR | **3.3** | 2.9 | 3 |
|  | 1E4Y | **2.6** | 2.4 | 166 |
|  | 1E4V | **2.8** | 2.5 | 153 |
|  | 2ECK | **2.8** | 2.6 | 143 |
|  | 1ANK | **2.9** | 2.6 | 136 |

The advantage of RRTMC compared to other methods of exploring protein conformational spaces can be summarized as being more time-efficient, being able to generate conformations that are close enough to the goal conformation (less than 3*Å*), and generating intermediate conformations that are close to experimentally known PDB conformations. (Refer to Table 1).

## 4 Conclusion

Exploring the conformational space of proteins is of primary importance for understanding their functions. Many proteins undergo large-scale conformational changes in order to perform their functionalities. Therefore, it is important to study these conformational pathways to find intermediate conformations that may correspond to local energy minima. It is experimentally difficult because these conformations are transitory and are hard to capture experimentally. To address this problem, computational methods are used to simulate these pathways. We developed a robotics-inspired algorithm using the RRT method and Monte Carlo, called RRTMC, to generate random conformations by rotating dihedral angles and added rigidity analysis information to make it run more efficiently and converge better. In this paper we presented an evaluation of our method using the Topological Data Analysis algorithm Mapper to find the intermediate conformations frequently produced by our algorithm, and compare them to experimentally available data. Our results show that TDA Mapper is highly beneficial to complement computational methods and that RRTMC is capable of generating conformations that approximate highly populated intermediate structures.

Current and future work includes using Mapper to evaluate multiple other proteins with available experimental data Running these experiments on more proteins is going to be our foremost important goal in future work. Another goal is exploring conformational transitions for protein in closer to real-life conditions. For example, the open conformation of AdK is dimeric in nature, and it would be interesting to explore the conformational transition using symmetry. Evaluating how Mapper works compared to other methods like K-means will be carried out. We plan to incorporate other dimensionality reduction methods such as Auto Encoders to compare the results. We also intend to use other clustering techniques instead of single-linkage in the Mapper algorithm to improve our algorithm.

## 5 Acknowledgment

This research was funded in part by NSF grant IIS:2031260. The tests were run on the supercomputing facilities managed by the Research Computing Department at the University of Massachusetts Boston, Chimera which is a heterogeneous distributed memory high performance compute cluster (AMD Opteron 6128 processors, 2.0 Ghz).

